# Spatial Confinement Promotes Hyphal Morphogenesis and Growth Advantage in *Candida albicans*

**DOI:** 10.64898/2026.11.26.769434

**Authors:** Lucie Albert, Antonio Serrano, Julie Foncy, Laurent Malaquin, Catherine Villard, Robert Arkowitz, Morgan Delarue

## Abstract

*Candida albicans* is an opportunistic human pathogen that can colonize multiple organs. During the commensal-to-pathogen transition, it often faces mechanical challenges and natural spatial confinement. Using molded agarose structures mimicking the geometry of an intestinal crypt and microfluidic confining devices, we demonstrate that spatial confinement induces a robust yeast-to-hyphal transition. Tip growth persists even after mechanical stress is removed. Furthermore, the hyphal form of *C. albicans* exhibits a growth advantage over the yeast form in mechanically constrained environments, likely due to reduced intracellular crowding. Our results suggest that mechanical cues modulate *C. albicans* morphogenesis, enabling escape from confined spaces.

## Article

The dimorphic nature of the human commensal *Candida albicans* enables it to colonize diverse host niches, from the gastrointestinal tract to mucosal surfaces^1,2^ , by switching between yeast and hyphal forms in response to environmental cues^3^. *C. albicans* can also grow as pseudohyphae, chains of elongated cells that remain attached after division and, unlike true hyphae, are constricted at the septal junctions^4^. This morphological plasticity is a hallmark of its adaptability, allowing it to transition from a harmless commensal to a pathogenic state under favorable conditions^5^. The yeast-to-hyphal transition is finely regulated by a combination of chemical and physical signals, including pH, temperature, nutrient availability, and quorum sensing molecules such as farnesol, which can either promote or inhibit filamentation depending on their concentration and context^3,6–9^. This transition is critical for *C. albicans* to evade immune responses^10^, invade tissues^11^, and form biofilms^12,13^, all of which contribute to its virulence and persistence in the host.

Superimposed on these physical and chemical cues, mechanical signals can also impact the virulence of *C. albicans*. Hyphal cells can produce significant forces^14^, and the microstructure of the environment can guide hyphae in a process called thigmotropism^15,16^. Cell growth in a confined environment, which can be observed during biofilm growth or cells trapped in small spaces like gut crypts under conditions of mucosal barrier deterioration^17,18^, can develop a growth-induced compressive stress^19–21^. In parallel, embedding cells in a solid agar matrix induces filamentous growth, a response first linked to the transcription factor Czf1^18^ and later to the Rac1/Dck1 GTPase module^22^. Yet in these embedded assays the physical stimulus is poorly controlled: matrix contact, mechanical resistance and local chemical gradients are confounded and forces were not measured, making it hard to assess the actual trigger of filamentation.

We 3D-printed molds to mimic crypt shapes (Fig. S1a) to test whether *C. albicans* proliferating in partial confinement could induce a yeast-to-hyphal transition. We seeded cells at 30°C, pH 5.5, in synthetic complete dextrose (SCD) medium onto these agarose-molded 400 µm height crypt-like structures of 50 µm in diameter at the bottom and 100 µm at the top (Fig. 1a). Note that this condition is not a full embedding but only a partial one, as the crypts remain open.

**Figure 1:**
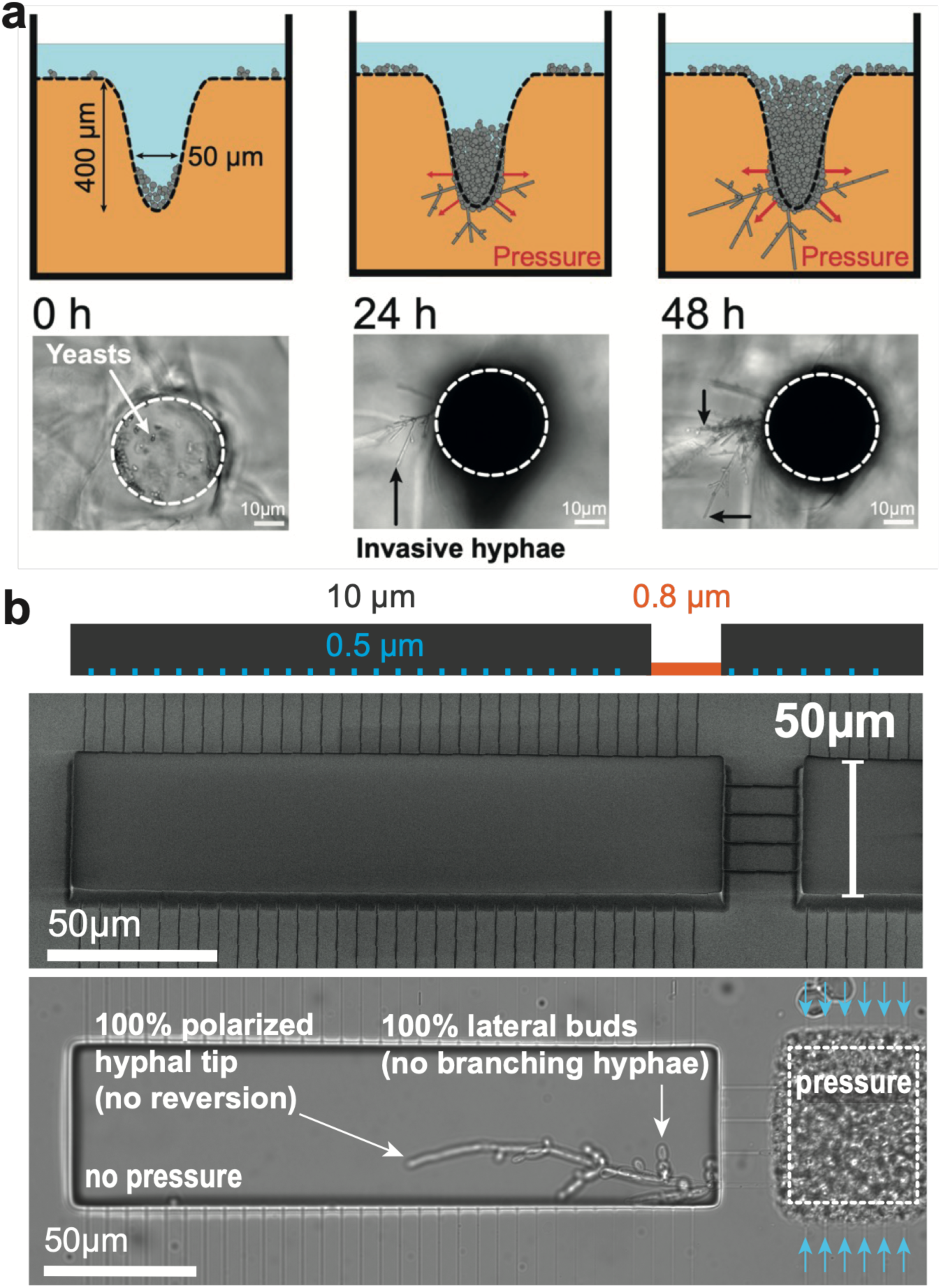
Partial confinement promotes a robust yeast-to-hyphal transition. **a.** *C. albicans* cells loaded onto a 2% w/v agarose mold reproducing the geometry of gut crypts developed a progressive growth-induced pressure. After 24 h of growth, hyphae invaded the surrounding agarose. Cells were cultured in synthetic complete dextrose medium at pH 5.5, 30°C. **b.** A 3-layer microfluidic device was developed to confine yeast cells and allow hyphae to invade into a relaxation chamber. After 5-6 h, growth-induced pressure was generated by the cells, and hyphae escaped the confining chamber to invade the relaxation chamber, where the tip cell grew as a hyphal cell for as long as we imaged. All the branched cells were in the yeast form. Cells were cultured in synthetic complete dextrose medium at pH 5.5, 30°C.

These culture conditions do not lead to hyphal morphogenesis in a liquid environment. After 24 h of partial confinement, we observed the appearance of hyphae, associated with enlargement of the crypt. We used finite-element simulations (Fig. S1b) to estimate the compressive stress generated from partial confinement to be on the order of a few kPa (Fig. S1c), which is in the physiological range. In this open geometry, however, there was no rinsing, and mechanical stress can be confounded with other chemical cues, such as changes in pH, oxygen, or even quorum sensing molecules concentration.

We then developed a 3-layer microfluidic device with a similar 50 µm width to promote spatial confinement (Fig. 1b), as in^19^. The chamber was connected on its sides to a set of narrow channels (0.5 µm × 1 µm) that were impassable for both yeasts and filaments. We connected this chamber to a relaxation chamber through a set of wider channels (1.5 µm × 1.5 µm), which were impassable for yeast cells but not for hyphae. When *C. albicans* cells grew confined at 30°C, pH 5.5, under a constant flow of SCD medium, they progressively built mechanical pressure (Video S1). We observed the growth of hyphae in the relaxation chamber. Interestingly, tip growth persisted in the hyphal form for as long as we could image them (more than 15 h). FRAP experiments between cellular compartments showed no chemical communication at the size scale of GFP (Fig. S2), ruling out potential protein-size cytoplasmic flow that would propagate from the mother cell in the confining chamber to the hyphal tip in the relaxation chamber. When new cells branched in the relaxation chamber, they branched as a yeast form, which progressively depolarized after a few generations (Fig. S3). Together, these results suggest that spatial confinement triggers a persistent filamentation in conditions that are usually not permissive for hyphal induction, but that this filamentation is not transmitted to daughter cells born outside confinement.

We used a fluorescent reporter of filamentous initiation and growth, *P_HWP1_-GFP*, to visualize and quantify when and where the switch occurred during confined growth (Fig. 2a, Video S2). We counted the number of inductions and reported them in space and time (Fig. 2b). Strikingly, we observed that the inductions were localized close to the inlet channels, and started to occur at very low mechanical pressure (below ∼ 15kPa, which is our minimal pressure resolution given a pixel size of 0.1 µm). Note that pressure can be heterogenous, but that there are no mechanical gradients in this setting^19^ : the induction gradient can only come from a chemical gradient. Stopping the advective flow, which would prevent the washing of quorum sensing (QS) molecules but did not prevent cell growth and pressure generation, had a dramatic effect on filament induction (Fig. S4a). In addition, increasing the inlet pressure from 1.2 bar to 3 bar increased advection inside the confining chamber and led to a larger gradient of induction (Fig. S4b). Finally, the addition of pravastatin at 64 µg/mL, which partly inhibits the QS farnesol molecule secretion^23^ , led to more induction (Fig. S4c). Taken together, our data suggest the existence of a spatial gradient of an inhibitory QS molecule, like farnesol^23^, superimposed on the mechanical compression induction signal.

**Figure 2:**
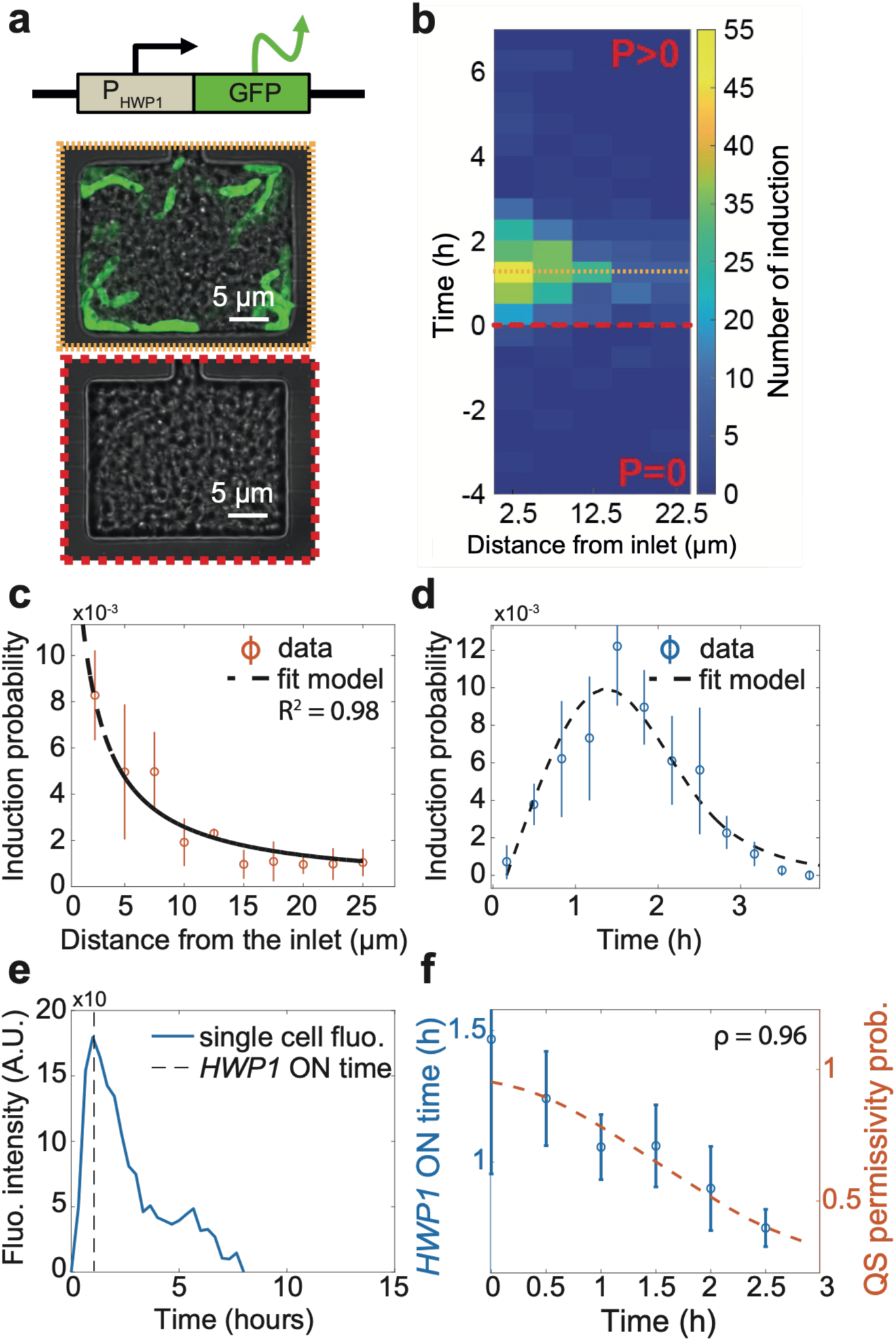
Spatial and temporal regulation of the yeast-to-hyphal transition under total confinement. **a.** We used a *P_HWP1_-GFP* fluorescent reporter of the yeast-to-hyphal transition. After confinement, cells expressing fluorescence could be observed in the confining chamber. **b.** We measured in time and space the occurrence of the yeast-to-hyphal switch, and observed that in SCD at pH 5.5, 30°C, the induction of hyphal cells occurred only after pressure was generated and was restricted to the sides of the confining chamber, and at early timescales. **c.** Probability of induction as a function of distance from the inlets for N ≥ 5 independent experiments containing at least 20 chambers in each replicate. Dashed line: fit of the data, R^2^ = 0.98. **d.** Probability of induction as a function of time after pressure generation for the same N ≥ 5 independent experiments containing at least 20 chambers in each replicate. Dashed line: fit of the data, R^2^ = 0.95, leading to ℙ_pressure_ = 0.095 ± 0.01. **e.** Example of the time evolution of fluorescence intensity at the single-cell level, displaying a maximum at about 1 h after fluorescent reporter induction. This maximum corresponds to the end of active transcription of the gene. **f.** Duration the reporter gene is active as a function of time, superimposed on the estimated permissive probability of quorum sensing molecules. The duration data aggregates 87 single-cell analyses over more than 5 independent replicates. ρ is the Pearson correlation result between the 2 datasets. In all experiments: mean ± SEM.

We developed a mathematical framework of advective transport inside a porous medium to model the quorum sensing concentration, *c*, gradient in the chamber (see Mathematical modeling section in SI). One implicit hypothesis is that the inhibitory QS molecules are secreted similarly for all cells, and in particular, irrespective of their position in the cell cycle. We found that we could define the quorum sensing permissivity probability, ℙ_QS_, from a first-order inhibitory function, ℙ_QS_ = ℙ_O_/ (1 + (*c*/*c*_O_)*^α^*), with *c*_O_ the inhibitory concentration and ℙ_O_ the probability at *c* = 0 accounting for pressure and cell cycle effects (see below). We estimated *α* = 1 from farnesol inhibition of a strong induction of hyphal transition (37°C, pH 7, serum, Fig. S5). From this function (see Mathematical modeling section in SI) we predicted a spatial dependence of hyphal induction to be in the form: 1/(1 + *x*/*x*_O_), which captures our data (Fig. 2c). From this, we could predict that faster flow washes QS inhibitory molecules, leading to the increased induction of hyphae when increasing inlet pressure to 3 bar (Fig. S4b).

We thus posit that the probability to filament in our setup, ℙ, is the product of three independent probabilities: ℙ = ℙ_pressure_ × ℙ_QS_ × ℙ_cycle_. ℙ_pressure_ would be the real probability to induce a filamentation under mechanical pressure. We assume it is null without pressure, and constant under pressure. The total probability to induce, ℙ, can be modulated by QS molecules, through ℙ_QS_, described above. Finally, ℙ could also be modulated by the probability of being in a given phase of the cell cycle, ℙ_cycle_. Indeed, in the chamber, cells were asynchronous, and although few reports suggest hyphal induction can happen anytime in the cell cycle^24–26^, we never observed budded cells expressing fluorescence from the *P_HWP1_-GFP* reporter. We observed a delay in the induction of hyphae, with a maximum induction of all hyphae in the chamber at about 1.5–2 h after confluence (Fig. 2d). This led us to hypothesize that only unbudded cells could morph into filaments in our conditions, and that this delay could represent a transition time through G1. We counted cell number in the chamber in time and measured that division rate decreased with pressure (Fig. S6a), similar to previously published work in *S. cerevisiae*^19,20^. This measurement allowed us to calibrate the probability to transit through G1 in an asynchronous population, ℙ_cycle_. We found that this probability decreased with time (Fig. S6b and Supplementary Information).

From the mathematical model in SI, we predict that the linear gradient of QS molecules increased with cell number (Fig. S6c). This allowed us to estimate the increase in overall concentration of QS molecules (Fig. S6c) from the increase in cell number over time (Fig. S6d), and to fully calibrate ℙ_QS_ as a function of pressure. Since ℙ_cycle_ was calibrated from the impact of pressure on cell division, we could fit the temporal evolution of hyphal induction (Fig. 2d), and extracted the mechanical probability, ℙ_pressure_, to be 0.095 ± 0.01. This means that, in constant chemical conditions, G1 cells have a 10% chance to become a filament under pressure. Interestingly, single-cell fluorescent data allowed us to extract the time the *HWP1* promoter was activated (Fig. 2e). We found that this ON time decreased with experimental time and followed the same trend as ℙ_QS_ (Fig. 2f). This is consistent with QS molecules modulating the activation time of filamentous genes under pressure.

We next sought to investigate the effect of spatial confinement on the growth and biosynthesis of both the yeast and the hyphal form of *C. albicans*. Similar to *S. cerevisiae*, we estimated the growth rate from the division rate, assuming balanced growth^20^ . We found that both growth rate and protein production from the *ENO1* promoter decreased exponentially with increasing compressive stress (Fig. 3a,b). We observed that hyphal extension was linear under pressure (Fig. 3c), and similar to the yeast growth rate measured at the population level, the extension rate decreased exponentially with mechanical stress (Fig. 3d), along with a decrease in GFP production from the *HWP1* promoter (Fig. 3e). Filament diameter seemed to remain constant with pressure. Interestingly, we observed that when normalized to the unconfined case, the pressure dependence of yeast and hyphal growth was not similar, hyphae being less sensitive to confinement than yeasts (Fig. 3f), suggesting that hyphal cells had an increased growth advantage under confinement.

**Figure 3:**
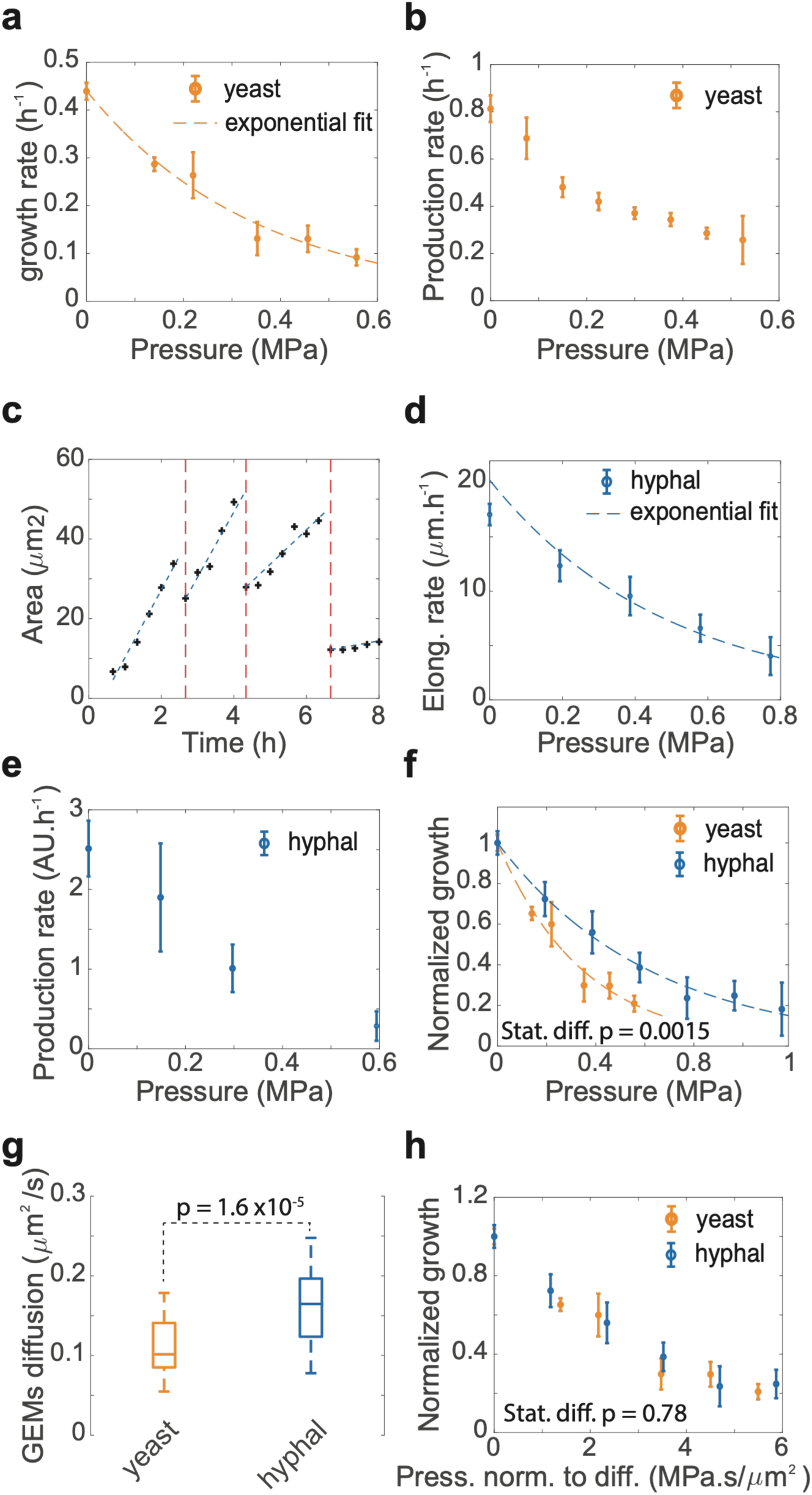
Spatial confinement slows both yeast and hyphal growth, but hyphae retain a growth advantage. **a.** Yeast growth rate as a function of growth-induced pressure. Dashed line: exponential fit. **b.** Production rate of *P_ENO1_-GFP* as a function of growth-induced pressure. **c.** Representative area of a growing hyphal cell as a function of time after confinement. Dashed line: linear fit to extract elongation rate. **d.** Hyphal elongation rate as a function of growth-induced pressure. Dashed line: exponential fit. **e.** Production rate of *P_HWP1_-GFP* as a function of growth-induced pressure. **f.** Normalized yeast growth rate and hyphae elongation rate as a function of growth-induced pressure. Statistical difference was assessed through a F-test, p = 0.0015. **g.** Diffusion coefficient of 40nm-GEMs in control yeast and hyphal cells. Statistical difference was estimated by a Mann-Whitney test, p = 0.000016. **h.** Normalized yeast growth rate and hyphal elongation rate as a function of growth-induced pressure normalized to 40nm-GEMs diffusion coefficient. Statistical difference was assessed through a F-test, p = 0.78 shows no statistical differences. In all experiments: mean ± SEM in at least 3 independent experiments, except for **a** and **b** (2 experiments).

It was shown in *S. cerevisiae* that macromolecular crowding, which relates to the multiscale active property of a biological milieu like the cytoplasm^27^, can limit confined cell growth^20^. Related to this result, we measured the diffusion coefficient of 40 nm genetically-encoded nanoparticles as a proxy for macromolecular crowding^28,29^, and found that hyphal cells were less crowded than their yeast counterparts (Fig. 3g). When pressure was rescaled by the unconfined diffusion coefficient (P = 0 MPa), both growth and extension rates of yeast and hyphal cells collapsed onto the same curve.

Together, our data demonstrate that spatial confinement and its associated growth-induced compressive stress can trigger a robust yeast-to-hyphal transition, even in conditions that are usually non-permissive for morphogenesis. Since cells are in contact with the chamber walls even before growth-generated pressure builds, the trigger is the kPa compressive stress itself rather than surface contact, although we cannot exclude that contact sensing can synergize with pressure. This transition is also persistent: tip growth persists more than 15 h in the relaxation chamber after stress removal. However, FRAP shows no protein-size cytoplasmic flow between compartments and new branches progressively revert to yeast, consistent with dilution of polarizing factors in daughter cells (Figs. S2 and S3). Spatial confinement occurs in physiological settings, from biofilms to gut crypts, and since kilopascal-level stress suffices, surface growth may be sensed by a similar mechanism^30,31^. Superimposed on the mechanical signal, chemical cues such as QS molecules can modulate the filamentous switch. We in particular find that higher flow decreases the inhibitory probability of QS molecules, making it more likely for filaments to form in catheters for instance.

Moreover, we find that hyphal cells have a growth advantage over yeast cells in mechanically constrained environments, which could allow them to better escape confined spaces and invade tissues. This advantage correlates with the lower intracellular crowding of hyphae (40-nm GEMs diffusion): yeast growth and hyphal elongation rates collapse onto a single curve when pressure is rescaled by the unconfined diffusion coefficient. This decreased crowding can impact many biochemical reactions^32^, making cells less sensitive to pressure^33^, but can also be associated with changes in turgor and / or cell wall properties.

While the molecular determinant of the mechanically-induced transition is elusive so far, surface mechanosensors such as Msb2^34^ or cell wall sensors^35^ are likely at play. Our results reframe earlier work on physically induced filamentation. Growth of cells embedded in a solid agar matrix has long been known to trigger filamentation, leading to the identification of Czf1 and, later, of the Rac1/Dck1 module as required for matrix-embedded invasive growth^18,22^. In those assays, however, the stimulus is binary and unmeasured, and matrix contact, mechanical resistance and chemical gradients are confounded, which precludes quantifying the mechanical / surface sensing contribution. Our calibrated, flow-controlled chambers show instead that a defined, self-generated compressive stress is sufficient to induce the transition under non-permissive chemistry. Whether this mechanical route converges on the Rac1/Dck1 axis remains an open question that our device can directly test.

## Acknowledgments

We thank Olivier Liot for useful discussions and proofreading of the mathematical modeling of advection in a porous medium. This work was supported by the LAAS-CNRS micro and nanotechnologies platform, a member of the Renatech French national network, the LAAS-CNRS characterization platform including MultiFAB platform, by an 80prime CNRS PhD grant (Lucie Albert) and by the ERC Starting Grant UnderPressure (grant agreement number 101039998). Views and opinions expressed are, however, those of the authors only and do not necessarily reflect those of the European Union or the European Research Council. Neither the European Union nor the granting authority can be held responsible for them.

## Author contribution

LA and MD designed and analyzed all the experiments with CV contributing as LA’s PhD co-advisor. LA performed all the experiments. AS and RA performed the GEMs experiments and analysis. JF and LA created the blueprint of the crypt-like structure, and JF 3D-printed it. LM performed the finite-element simulations. MD and CV financially supported the work. MD and LA wrote the manuscript, and all the authors proofread it.

## Data availability

All the data, code and blueprints will be deposited on a public repository once the article is published.

## Material and Methods

### Strains used and cell culture

Cells were thawed routinely from -80°C stock every two weeks and grown on YEPD agar 65g.L-1 (Sigma-Aldrich) at room temperature. Strains were serial diluted in buffered liquid Synthetic Complete medium (SC) with 400mg.L-1 Complete Supplement Mixture (Formedium), 750mg.L-1 Yeast Nitrogen Base (Sigma-Aldrich) and 20 g.L^-1^ dextrose (Sigma-Aldrich) buffered at pH5.5 (10 mM monohydrate MES, Sigma-Aldrich) and grown overnight at 30°C and 200 rpm. Suspensions at an optical density (OD_590nm_) between 0.1 and 0.3 were selected before live-imaging experiments. Unless otherwise indicated, the same-buffered SCD culture medium was used during all live-experiments.

The *Candida albicans* strain (PY6414) expressing 40-nm genetically encoded multimeric nanoparticles (GEMs) has been described previously^29^. For budding growth, cells were cultured at 30 °C in YEPD supplemented with uridine (80 µg ml⁻¹). Filamentation was induced by transferring cells to YEPD containing 50% fetal bovine serum (FBS; PAN Biotech) and incubating them at 37 °C for 45–150 min. Budding cells were shifted to 37 °C shortly before imaging, so that both cell types were imaged at the same temperature.

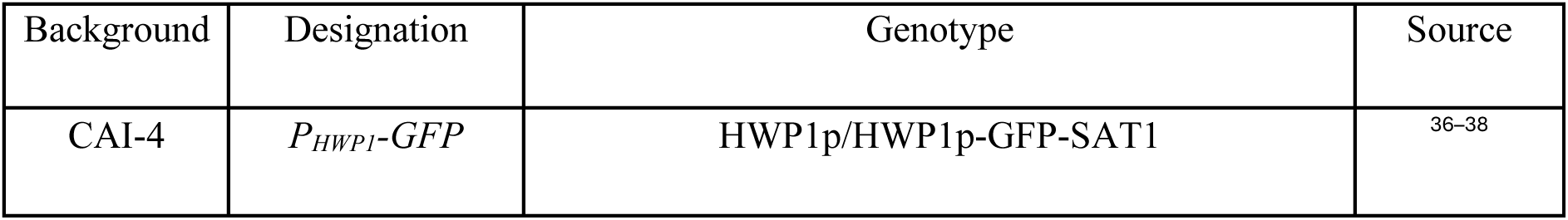
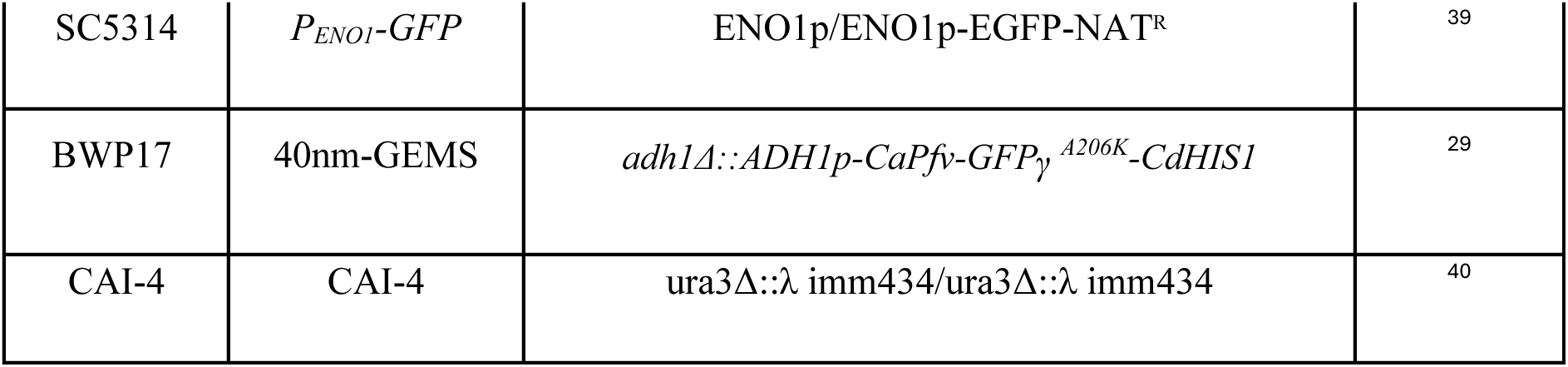

| Background | Designation | Genotype | Source |
|---|---|---|---|
| CAI-4 | *P<sub>HWP1</sub>-GFP* | HWP1p/HWP1p-GFP-SAT1 | <sup>36–38</sup> |
| SC5314 | *P<sub>ENO1</sub>-GFP* | ENO1p/ENO1p-EGFP-NAT<sup>R</sup> | <sup>39</sup> |
| BWP17 | 40nm-GEMS | *adh1Δ::ADH1p-CaPfv-GFPγ<sup>A206K</sup>-CdHIS1* | <sup>29</sup> |
| CAI-4 | CAI-4 | ura3Δ::λ imm434/ura3Δ::λ imm434 | <sup>40</sup> |

### Imaging conditions

Partial cell confinement in crypt-like structure assay was performed on a Leica DMi8 epifluorescence inverted microscope equipped with a motorized stage, an automatic focus, and a scMOS camera (Hamamatsu, ORCA-Flash4.0). Image acquisition was achieved using a 40x immersion objective (Leica, 17/E PL AP0 1.25/0.75 NA oil) and a LED illumination unit (Lumencor, Spectra Light Engine). The microscope and its components were operated by Micromanager 2.0 software. The microfluidic device was placed in a thermo-regulated chamber (Tokay Hit, STX) set to 30°C.

Mean volume fraction of cells in the confinement compartments was measured by exclusion of fluorescence as in^19^ by adding FITC-DEAE-Dextran (Sigma-Aldrich MW 3,000-6,000) at 40 µM to circulating SCD at the beginning of the live-cell experiment. LED intensity was set to 10% of nominal power (GFP, bandpass filter 542/30m, nominal power 400 mW). Images were acquired every 20 min in GFP and every 10 min in bright field to assess corresponding deformation of the confinement compartments.

Fluorescence Recovery after Photobleaching (FRAP) was performed on a Zeiss 710 inverted confocal microscope equipped with a thermo-regulated chamber operated using Zen Blue software. Acquisitions were performed using a GFP filter set (Excitation BP470/40 DM495 Emission BP525/50 and GaAs BP500-550 nm/GaAs). Laser argon (488 nm) was set to 50 % of nominal power and exposure to 400ms for bleaching. Images of the bleached ROI were acquired at 10 s interval over a 60s total period using a 100x oil immersion objective (Leica, PL APO oil DIC 1.4 NA) and bleaching was performed after the first image.

Time-lapse microscopy experiments were performed on a Leica IRB microscope equipped with a confocal spinning-disk unit (Yokogawa, CSU-X1, nominal power 100 mW), a motorized stage, an automatic focus, and a scMOS camera (Hamamatsu, ORCA-Flash 4.0). Either brightfield or fluorescence images were acquired with an inverted 63x oil-immersion objective (Leica, HC-PL APO 63X 1.40/0.60 NA). Fluorescence signaling was acquired with appropriate filter set, either ET525/50m (GFP), ET595/50m (RFP) or ET450/50m (DAPI) combined with a dichroic mirror (Chroma, ZT405/488/561/640rpc). The microscope and its components were operated using Micromanager 1.4 software. The fluorescence signal derived from either pHWP1-GFP-SAT1 reporter of hyphal growth or pENO1-GFP-SAT1 reporter was monitored overnight every 20min in GFP. For assessing cell number at the bottom plane (near glass coverslip) RhodamineB-Dextran (Sigma Aldrich, 10,000 MW) was added to culture medium at 20µM. Fluorescence signalling was monitored every 60min in RFP. GEMs images were acquired at a framerate of 100Hz (period 10ms) under continuous exposure (laser 488nm). Particle tracking was achieved using ImageJ MosaicSuite (v. 1.0.24) and effective diffusion was inferred with a home-made MATLAB (v. 2020b) script.

Bright field images were acquired as well every 10min to assess lateral deformation of confinement compartments and infer growth-induced pressure.

### Crypt-like mold printing and agarose molding

Crypt-like mold was designed with FreeCad as a cylindrical stamp (2.45 mm thickness, 5.0 mm diameter) comprising spacers (thickness 0.45mm) and micrometric pillars on one side. Conical micropillars were 0.44mm high with diameter of 0.05mm at the tip and 0.1 at the bottom.

Molds were fabricated via stereolithography using a Dilase Bio 3D printer (Kloé SA, France) equipped with a 405 nm laser source and an optical objective yielding a 10μm beam diameter. 3D CAD model was sliced with proprietary slicing software (Kloé SA, France) at a 10μm slice thickness. The base of the mold was printed with an exposure dose of 10.02 mJ/cm2 (99% laser power, 80mm/s scan speed, and 2 mm of stabilization), while the pillars were printed using a dose of 0.51 mJ/cm2 (2% laser power, 31 mm/s, scan speed, and 0.5 mm of stabilization). Printing time 5.5 h. After printing, the printed mold was detached from the building platform, rinsed with ethanol, and developed in an ultrasonic bath (P-Series Ultrasonic cleaner, Fisherbrand™) for 3 minutes. The pieces were rinsed a second time with ethanol, dried, and subjected to post-polymerization in a UV chamber (32 mW) for 30 minutes.

20 µL buffered SCD with 2% w/v agarose was heated then molded at room temperature in µ-Plate 24 well with polymer coverslip bottom 180µm (Ibidi) by gently pressing the mold for one minute. Resulting agarose structure was then covered with 1mL of cell suspension from fresh overnight culture in buffered SCD medium and was kept static for 2 hours at 30°C to allow the sedimentation of cells into crypt-like structures. Excess liquid was gently flushed and rinsed with 2mL of fresh buffered SCD medium and kept under agitation at 30°C overnight. Overnight culture was gently rinsed with 2mL of fresh SCD medium to remove excess cells above the structure and observed

### Finite-element simulations to estimate generated pressure in molded crypts

Finite element simulations were performed using COMSOL Multiphysics to investigate the mechanical deformation of hydrogel-based crypt structures subjected to internal pressure. The agarose hydrogel was modeled as a homogeneous, isotropic, linear elastic material. Crypts were represented using a 2D axisymmetric geometry, with a depth of 400 µm, an opening diameter of 50 µm, and a rounded bottom with a radius of curvature of 50 µm. The bottom surface, corresponding to the glass–hydrogel interface, and the external lateral boundaries were fixed.

The exact mechanical properties of agarose are hard to extract, with the literature displaying many different values. We fixed the Young’s modulus to be 50kPa as an average value for agarose 2% measured in compression^36–38^ , and tested low (0.17) and close to incompressible (0.48) Poisson ratio (Fig. S1c). A uniform pressure ranging from 0 to 10 kPa was applied to the internal crypt surface. The resulting radial displacement of the crypt walls was evaluated at a depth of 100 µm.

The simulations revealed a linear relationship between the applied pressure and the lateral displacement of the crypt walls. For a representative Young’s modulus of 50 kPa, an applied pressure of approximately 4.8 kPa was associated with a lateral displacement of approximately 3 µm. The choice of the Poisson ratio merely impacted the estimated displacement.

### Microfluidic device mold preparation

Microfluidic device molds were fabricated as detailed in^41^. Briefly, photoresist master molds designed using Clewin5 were fabricated by standard photolithography process on silicon wafer. A first layer of ∼0.5µm height negative photoresist (Microchem, SU-8 TF 6000.5) was constructed to generate nano-inlets supplying nutrition and retaining cells. Microfluidic devices involving a mechanical relaxation compartment comprised a second ∼0.8µm height layer (Microchem, SU-8 2000.5) to form channels linking each compartment. A last ∼10µm height layer (Microchem, SU-8 3005) was fabricated to generate top structures and compartments. Photoresist molds were all silanized by vapor deposition of trichlorosilane to limit attachment from polydimethylsiloxane (PDMS).

The surface of mold was characterized by Scanning Electron Microscopy (MEB Hitachi S-4800). Tension and current were respectively set at 0.6 kV and 8 μA. To correct astigmatism, magnification was set at ×3000.

### PDMS chip preparation and usage

PDMS (DowCorning, Sylgard-184) were prepared from mixing curing agent and base at 1:10(w/w) ratio then cured onto master molds at 60°C overnight. PDMS chips were cut off the mold and a biopsy puncher (Robbins Instruments, OD 0.75mm) was used to open inlets for cell loading and medium circulation. Then, PDMS chips were cleaned with isopropanol, covalently bounded onto glass coverslips (24x60x0.17 mm) by plasma surface activation (Diener PICO, PO2 at 0.3 mbar, 50 W, 30 s) and baked at 60°C overnight. Freshly fabricated devices were used within less than a week to avoid PDMS nano-inlets collapse on glass.

Loading the microfluidic devices involves three main steps as described in^41^ . Yeasts were loaded into the confinement compartments using a syringe. Then, SCD was flushed through the circulating inlet to evacuate the air of the micro-channels *via* an outlet. Lastly, the outlet was sealed thus enabling SCD circulation through the confinement compartments containing yeasts during live-cell experiments. All microfluidic devices were treated with plasma (P_O2_ at 0.3 mbar, 50 W, 40 s) before experiments to increase channel hydrophilicity. Unless otherwise indicated, the working pressure applied to circulating SCD was set to 1.2bar (17.4psi) using a MFCS Fluigent.

When indicated, pravastatin sodium (MedChemExpress) a 3-hydroxy-3-methylglutaryl-CoA (HMG-CoA) reductase inhibitor was added to the perfused SCD at a working concentration of 64 mg.L^-1^ from a stock solution at 10 g.L^-1^ (diluted in ddH_2_O).

### Image analysis

#### Inferring growth-induced pressure (GIP) from confinement compartment deformation

Deformation of PDMS confinement compartments were measured from brightfield microscopy using a homemade semi-automated algorithm (Matlab, R2019b). Briefly, the script detects the edges of the compartment allowing measuring its width over time. Then, width values were converted to growth-induced pressure (GIP) values using a calibration curve constructed as in^19^ leading to a conversion factor of 7.5 µm.MPa^-1^.

#### Number of proliferating cells at the bottom

Total number of cells was evaluated using fluorescence exclusion from spinning-disk confocal microscopy. Cells at the focal plane were manually counted every hour using ImageJ (v.1.53) starting from inoculation and averaged over confinement compartments (two independent experiments). Only the first layer near glass coverslip was considered and cell density was estimated homogeneous throughout the XY plane.

#### Counting yeast-to-hypha transition occurrences in the confinement compartments

Time of appearance and spatial location of newly pHWP1-GFP cells expressing fluorescence were manually annotated using ImageJ (v.1.53). Each measurement was made within a 50x60 µm ROI centered on the confinement compartment to normalize all positions of inductions across images. The time at which cell population reaches confluency was manually estimated from bright-field images for each confinement compartment. Only primo-inductions were considered and cells that inherited fluorescence signal from parental yeast were not considered. Time of each primo-induction was shifted by subtracting the time at confluency. Location of occurrences relative to the nutrient nano-channels was normalized leading to values between 0 µm (near the nano-channels) and 25 µm (half-width of confinement compartment).

#### Estimating spatio-temporal yeast-to-hypha probability ℙ

Time and location of primo-inductions extracted from each confinement compartment were pooled together for each experiment. Location values were discretized and cumulative count of primo-inductions over time was calculated for different distances from nano-inlets, from 2.5 to 25 µm. Step-size between values was set to 2.5 µm and the width of slices to 2.5 µm. ℙ was inferred by dividing by the total number of proliferating cells contained in each.

#### Inferring mean volume fraction of cell in the confinement compartment

Mean volume fraction of cells was inferred from FITC-Dextran fluorescence intensity values as detailed in^19^ using the Beer-Lambert law. Denoting the packing fraction of cell, the volume occupied by the cell biomass, and the free volume in the confinement compartment. Fluorescence signal from the dye surrounding the cells was assumed proportional to the packing fraction of cell. To obtain a continuous function from experimental data, values were plotted as function of growth-induced pressure values for each confinement compartment and fitted by a second order exponential function. Cell volume fraction was estimated at confluency when growth-induced pressure begins to increase. Finally, porosity ε was inferred from the relationship ε = 1 − Φ.

### Single particle imaging and tracking

GEMs were imaged by total internal reflection fluorescence (TIRF) microscopy on an inverted Nikon Ti Eclipse microscope (Nikon France S.A.S.). The microscope was fitted with an iLas² scanning head (Roper Scientific), a ×100 CFI Apo TIRF oil-immersion objective (NA 1.49) and an iXon 888 EMCCD camera (Andor Technology). GFP was excited with a 488-nm diode laser at 30% power and a TIRF angle of 56.24°. For each field, 300 consecutive frames were recorded in a single focal plane at 30 ms per frame. Samples were held at 37 °C with an Okolab incubator during acquisition, and multiple fields per sample were collected using a motorized XYZ stage. Particle trajectories were obtained with the MOSAIC particle-tracking plugin^42^ in Fiji (v1.54f) running on a Windows 10. Effective diffusion coefficients (Deff) were then calculated in MATLAB (R2023a) as previously described^28^ . Filament length was measured in Fiji using the segmented line tool, traced along the hypha from its point of emergence at the mother cell to the hyphal tip.

### Statistics

A Pearson correlation was used to show the correlation of the ON time of HWP1 gene induction and the estimated permissivity probability of QS molecules (ℙ**_QS_**).

A Mann-Whitney U-test was used to test for the statistical difference in the diffusion coefficients of yeasts and hyphae in Fig. 3g.

To test whether the pressure dependence of yeast growth and hyphal elongation differed, we compared a single exponential decay, y = exp(−P/λ), fitted to all pooled points, against two separate exponentials fitted to each form separately. Curves were compared with an extra sum-of-squares F test (F test for nested models; equivalent to a likelihood-ratio test). We calculated F = ((RSS_shared_ − RSS_sep_)/(2-1)) / (RSS_sep_/(N − 2)), where RSS is the residual sum of squares of the shared (RSS_shared_) and separate (RSS_sep_ = RSS_yeast_ + RSS_hypha_) fits and N = 19 is the total number of points. The hypothesis of a common decay curve in Fig. 3f was rejected (F(1, N−2) = 14.21, p = 0.0015). When pressure was instead rescaled by each form’s unconfined 40-nm GEM diffusivity (Fig. 3h), the same test could not reject a common curve (F = 0.074, p = 0.78), showing that yeast and hyphal growth collapse onto a single pressure–growth relationship once intracellular crowding is accounted for.

**<u>Video S1</u>: Confined yeasts movie.** Confined yeasts morph into filaments that escape the confining chamber into a relaxation chamber. The tip cell persists as a filament for as long as we can image, while branching occurs as yeasts. Scale bar: 10 µm.

**<u>Video S2</u>: PHWP1-GFP construct.** Strong (pH 7, 37°C, serum) induction leads to filamentation, and accumulation of fluorescence in filaments.

---

**Figure S1:**
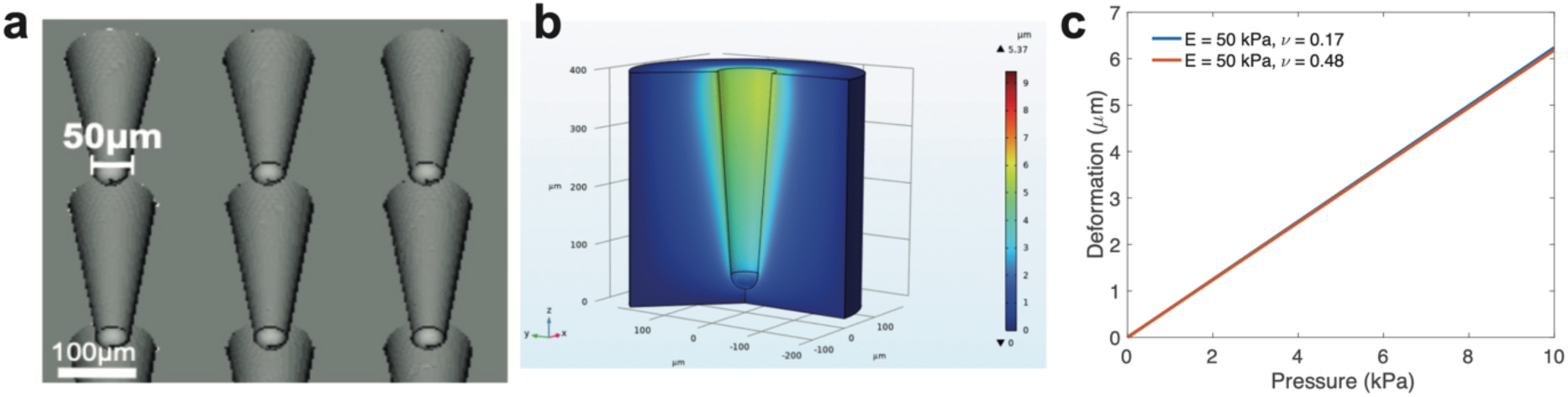
Mold and Finite-element simulation. **a.** Image of the CAD file used to 3D-print the crypt-like structures. **b.** Finite element simulation of the deformation of our crypt-like structure showing the Van Mises stress on the structure for a given displacement.

**Figure S2:**
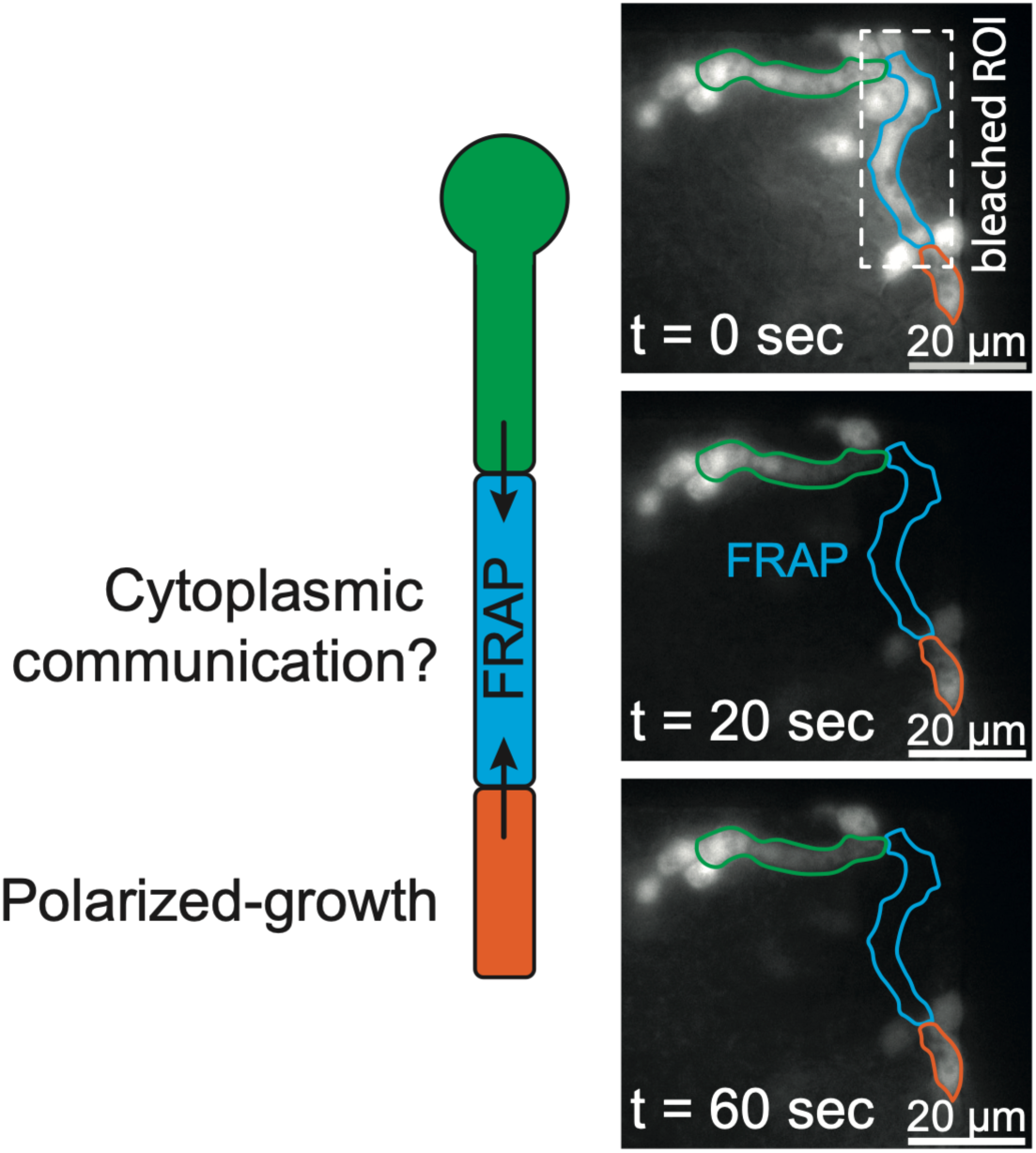
Photobleaching of a compressed yeast compartment. A compartment of a growing confined filament expressing GFP from the *HWP1* promoter was bleached. We observed no recovery of fluorescence from the adjacent compartments suggesting no protein communication under these mechanical conditions.

**Figure S3:**
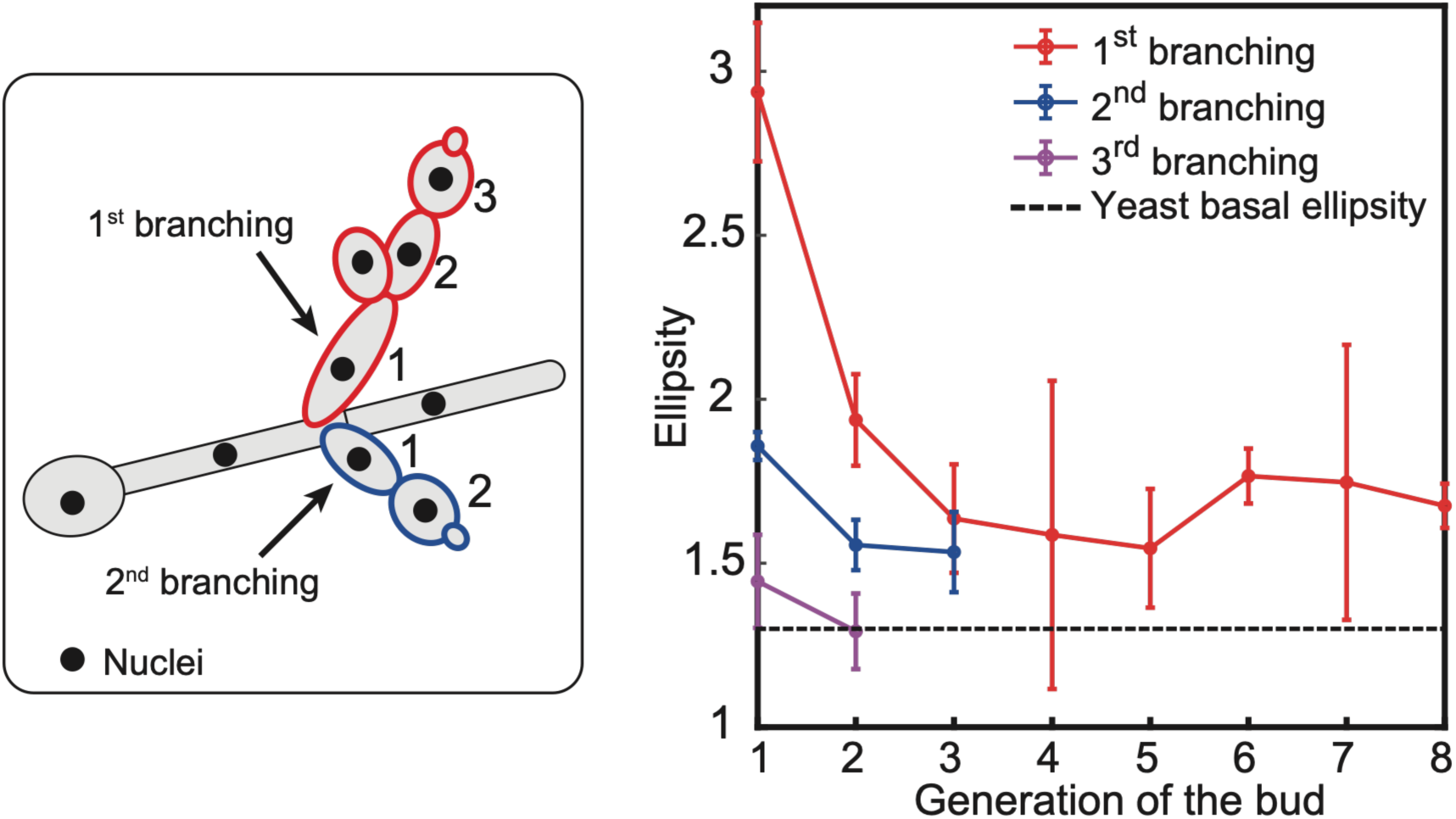
Elongation recovery in unconfined hyphae. Unconfined hyphae branched as yeast cells. We observed that these cells were less and less polarized as they divided, and that the third generation of branching had the same shape as the yeast form, suggesting a progressive dilution of polarizing factors as cells grow.

**Figure S4:**
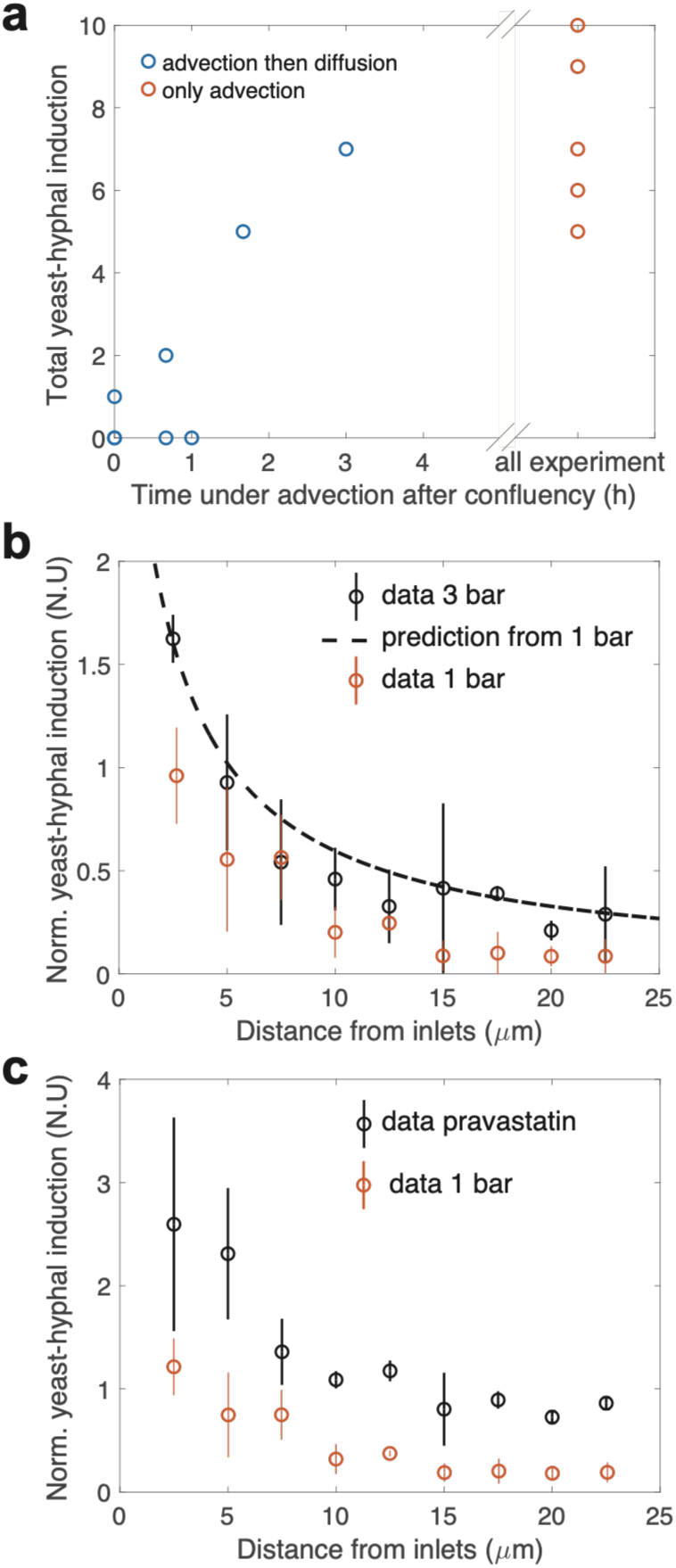
Inhibitory quorum sensing molecules create a gradient of hyphae under confined growth. **a.** We stopped the inlet flow of medium sometime after confluency, so that only diffusion prevails. We observed that if diffusion occurs right before or close to the onset of pressure generation, no hyphae are formed. Each point is a different chamber. **b.** Spatial probability of hyphae in a confining chamber with a higher flow rate (3 bars instead of 1.2 bars). The probability is normalized at 2.5 µm to the value of the 1.2 bar flow rate (control). **c.** Spatial probability of hyphae in a confining chamber with 64 µg/mL of pravastatin. The probability is normalized at 2.5 µm to the value of the 1.2 bar flow rate (control). For b and c, at least 3 independent experiments were performed and at least 5 different chambers were analyzed. Data represents mean ± SEM.

**Figure S5:**
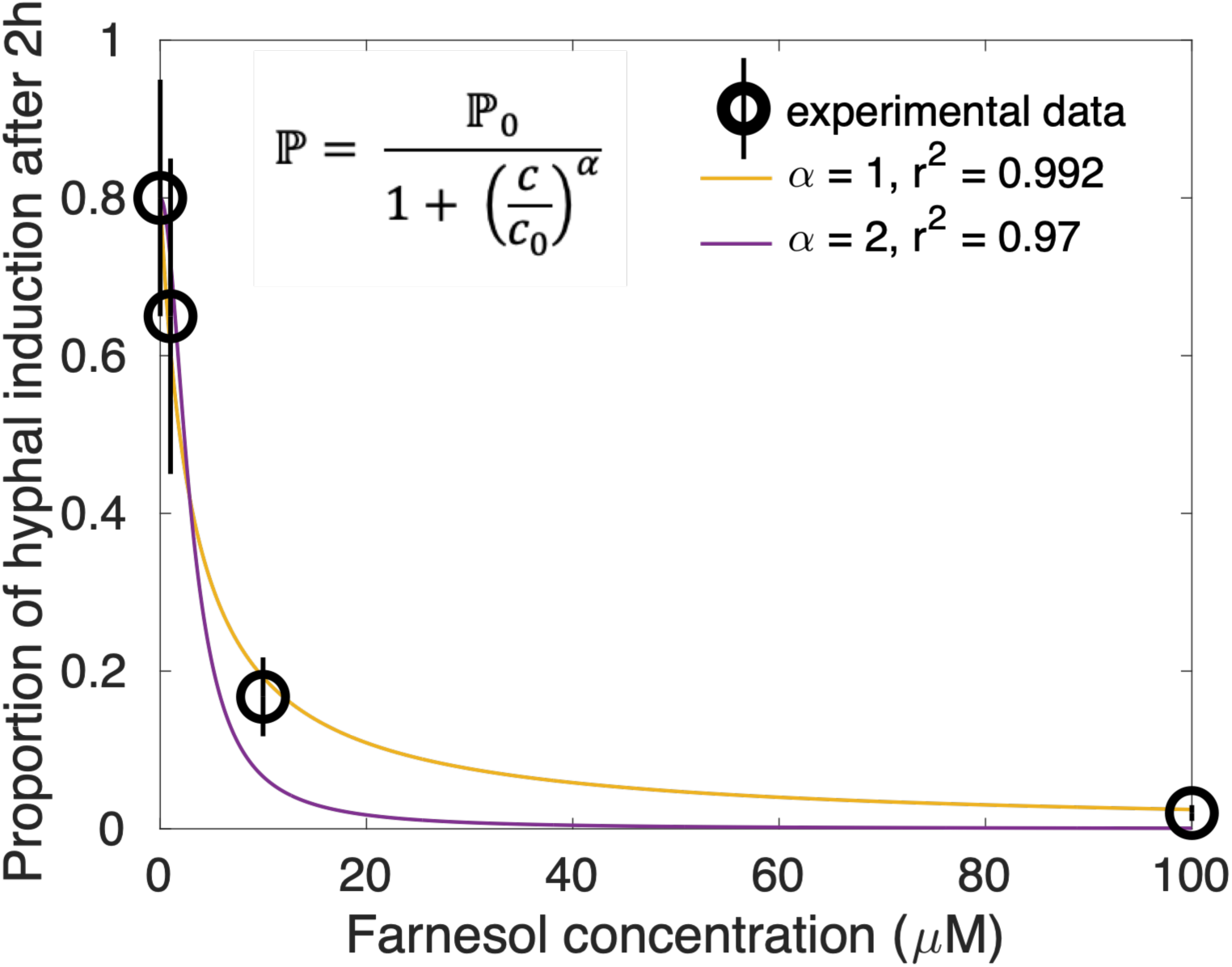
Probability of inducing hyphae under strong transforming conditions and farnesol. Cells were exposed to 37°C, pH 7 and a medium containing serum and farnesol. We measured the probability of cells to express GFP from the P_HWP1_-GFP construct 2h after induction. We fitted the data with the function displayed as an inset, and empirically observed that α = 1 is the best fit.

**Figure S6:**
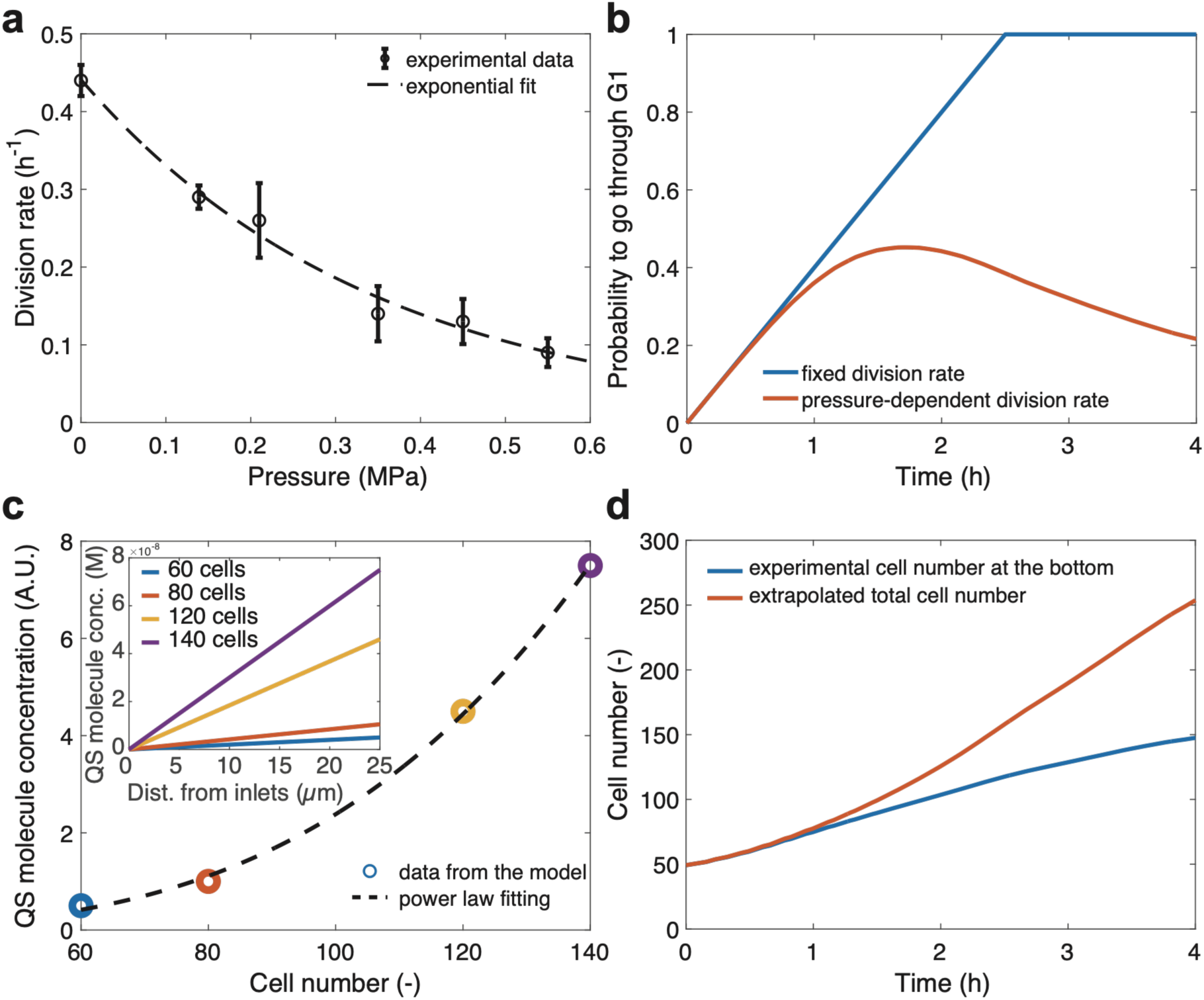
Confined growth impacts division rate and probability to go through G1, as well as inhibitory quorum sensing molecules. **a.** Cell division rate decreases exponentially with pressure. Division rate was measured from the derivative of cell number. At least 2 independent experiments were used to compute this metric. The decrease can be fitted by an exponential function. **b.** Probability of a cell to go through G1 if (i) the division rate is constant or if (ii) the division rate decreases with generated pressure. In the second case, since division rate decreases, the probability decreases at long timescale. **c.** Estimated of quorum sensing molecules at the center of the chamber as a function of cell number. These data can be fitted by a power law function. Inset: spatial gradient of quorum sensing molecules as a function of cell number. **d.** Measurement of cell number at the bottom of the chamber (only the mean is represented), and extrapolation to the total cell number in the whole chamber.

