## Supplementary Information for "Spatial Confinement Promotes Hyphal Morphogenesis and Growth Advantage in *Candida albicans*"

### Analytical model describing spatial gradient of quorum sensing molecules in the chamber

#### Preamble

We propose that yeast-to-hyphal probability is mainly modulated by three phenomena in this defined physicochemical microenvironment: (i) the fact that there is mechanical compression, (ii) the fact that they must go through the G1 phase of the cell cycle, and (iii) by the presence of quorum sensing molecules. Assuming that the three probabilities are independent, they are multiplicative:

$$\mathbb{P} = \mathbb{P}_{\text{pressure}} \times \mathbb{P}_{\text{cycle}} \times \mathbb{P}_{\text{QS}}$$

(i) We assume that $\mathbb{P}_{\text{pressure}}$ is a function that is 0 without pressure and has a constant value $\mathbb{P}_{\text{pressure}}$ when the pressure builds up. This value is a fitting parameter, discussed in the manuscript.

(ii) As we never observed budding cells displaying fluorescence from the $P_{HWP1}$-GFP reporter, we assume that only cells in G1 can transform. Under constant division time, the probability to be in G1 should increase linearly as $k_{\text{div}} \times t$ then saturate to 1 (Fig. S6b). However, as the division rate decreases with increasing compression, this probability decreases as $k_{\text{div}}(P) \times t$ decreases for large pressure $P$ (Fig. S6b). There is no fitting parameter here, as the numbers are calculated experimentally.

(iii) As shown in Fig. S4, there is a spatial gradient of some inhibitory quorum sensing molecules, likely farnesol. This modulates the probability to induce. In the following, we describe our reasoning for the calculation and calibration of $\mathbb{P}_{\text{QS}}$.

### Calculation of $\mathbb{P}_{\text{QS}}$

We observed that proportion of hyphae growing under inducible conditions (37°C, SCD + 20% serum) decreases with the exogenous farnesol concentration (Fig. S5). Especially, after 2h, the proportion of hyphae under increasing farnesol dose can be fitted with a first order sigmoid. Since we cannot know if there are other quorum sensing inhibitory molecule, we will more generally refer to QS molecule rather than farnesol. We posit that $\mathbb{P}_{QS}$, the probability of yeast-to-hyphal transition as function of quorum sensing molecule concentration, can be described as:

$$\mathbb{P}_{QS}(t) = \frac{1}{1 + \frac{C(t)}{C_{50}}}$$

where, C is the concentration of QS molecules secreted by the proliferating cells ($mol.L^{-1}$), and $C_{50}$ is the concentration level that represses 50 % of the potential yeast-to-hypha transitions under inducible conditions. In the following, we describe a mathematical solution of the concentration of QS molecules in 1D, along a flow line from the inlet to the center of the chamber, in the porous material that the cells create[1].

**1-D advection-diffusion modeling of farnesol transport**

To estimate the concentration of secreted QS molecules along the half-width of the confinement compartment $C(x, t)$, we developed an analytical model of advection-diffusion in a porous medium. We describe the concentration of a diluted specie C ($mol.L^{-1}$) in a porous medium starting from the law of mass conservation:

$$\varepsilon \frac{\partial C}{\partial t} = \varepsilon \operatorname{div}\left(\overline{\overline{D}} \cdot \overrightarrow{\operatorname{grad}} C\right) - \operatorname{div}(\langle \vec{v} \rangle \cdot C) + \Lambda \qquad (1)$$

where C is the intrinsic average concentration of QS molecule ($mol.L^{-1}$) over a representative elementary volume; $\langle \vec{v} \rangle$ is the superficial average velocity in the porous medium ($m.s^{-1}$); $\overline{\overline{D}}$ is the effective diffusion-dispersion tensor; and $\Lambda$ is the source term of the specie (effective farnesol secretion). Finally, $\varepsilon$ is the average porosity inferred from exclusion of fluorescence measurements (see Material and Methods). Experimental data and adjusted continuous function to estimate $\varepsilon(P)$ are displayed Fig S7:

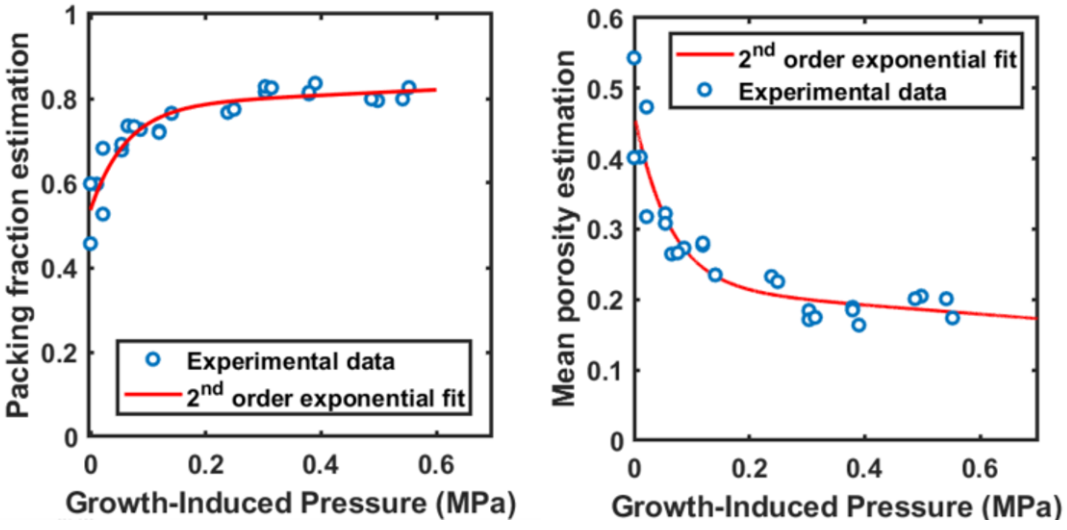


**Figure S7**: (left) Cell packing fraction ϕ increases with pressure. Experimental data were fitted with a 2nd order exponential. (right) Average porosity of the cell proliferating under pressure was calculated as $\varepsilon = 1 - \phi$.

Simplifying to 1D, equation (1) can be re-written:

$$\varepsilon \frac{\partial C}{\partial t} = \varepsilon D_{eff} \frac{\partial^2 C}{\partial x^2} - \langle \vec{v} \rangle_x \frac{\partial C}{\partial x} + \Lambda \qquad (2)$$

where, $D_{eff}$ defines the effective diffusion in the porous medium ($m^2.s^{-1}$); $\langle \vec{v} \rangle_x$ is the superficial average velocity along the considered abscissa (half-width of the chamber, detailed in Fig. S8); C is the intrinsic average concentration of farnesol ($mol.L^{-1}$) over a representative elementary volume and $x$ is the curvilinear distance relative to the feeding inlets along the half-width abscissa (Fig. S8). To simplify the problem, we approximated the curvilinear abscissa (streamline of the flow in the confinement compartment) as a linear abscissa along half-width of the compartment, parallel to direction of the nano-inlets.

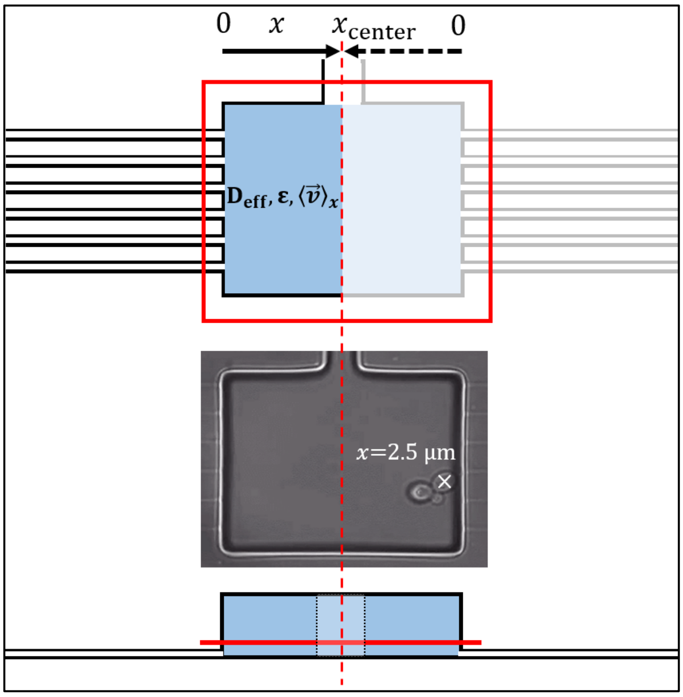


**Figure S8**: Picture and schematic view of the confinement compartment, and the considered porous medium. Rectangle in light blue represents the volume occupied by the porous medium (proliferating confined cells) with the characterized transport parameters (effective diffusion of solute, porosity, average velocity respectively). Red rectangle marks the ROI of 50×60 µm centered over each compartment to annotate position of Y-H occurrences. Dashed line highlights the compartment axial symmetry, as 6 nano-inlets supply medium on each side. An example of marked cell position in contact with compartment wall at nano-inlets is given (x = 2.5 µm by symmetry). Schematic of a chamber section at the bottom shows the position of the focal plane near glass-coverslip and nano-inlets (red plain line).

To describe QS molecule secretion, we assumed each yeast produces and adsorbs molecules at a constant rate ($s_{eff} = s_{production} - s_{adsorption}$; mol.$s^{-1}$), independently of mechanical pressure and cell cycle state. As we aim to describe a concentration, we divided this quantity by the elementary medium volume ($V_{elementary}$; $m^3$) surrounding each cell that we defined as follow:

$$V_{elementary} = \frac{\varepsilon V_{total}}{N_{total}} \quad (3)$$

where, $V_{total}$ is a volume of $5 \times 40 \times 50$ µm$^3$ containing the proliferating cell monolayer against the glass coverslip characterized by a porosity ε. Therefore, the quantity $\varepsilon . V_{total}$ represents the total free volume surrounding the observed cells. We express the local QS molecule secretion as follow:

$$\Lambda = \frac{s_{eff}}{V_{elementary}} = \frac{s_{eff} N_{total}}{\varepsilon V_{total}} \qquad (4)$$

We estimate the characteristic diffusion time τ in the porous medium using the following equation:

$$\tau = \frac{l_\tau^{\,2}}{D_{mol}^{eff}} \qquad (5)$$

where $l_\tau$ is the typical travelled length within the confinement compartment, and $D_{mol}^{eff} = \frac{D_{mol}}{T}$ the molecular diffusion coefficient of QS molecule corrected by the tortuosity of the porous medium T. It is hard to know the exact value of the tortuosity, but it can be bounded by ~ 2.15 (Ref [2]) for a low porosity (0.1), not affecting the molecular diffusion coefficient too much.

As an upper-limit value, we assume that $l_\tau \leq w_c$, where $w_c$ is the total width of the confinement compartment ($w_c = 50$ µm).

To our knowledge, the molecular diffusion of QS molecules such as farnesol has not been characterized yet. Thus, we approximated $D_{mol}$ with the diffusion of glucose in water at 30°C (740 µm$^2$.s$^{-1}$)[3] , a specie with a molecular weight similar to farnesol.

Considering these values, the typical free diffusion time of QS molecules is of the order of minute. This value is negligible compared to the typical cell proliferation time, as yeast cell-cycle span is about 1 hour. Therefore, we assume a near-stationary regime and neglect the quantity $\frac{\partial C}{\partial t}$. Finally, by introducing the expression of Λ, Eq. 2 simplifies as follow:

$$\varepsilon D_{eff} \frac{\partial^2 C}{\partial x^2} - \langle \vec{v} \rangle_x \frac{\partial C}{\partial x} + \frac{s_{eff} N_{total}}{\varepsilon V_{total}} = 0 \qquad (6)$$

##### Boundary conditions and analytical expression of QS molecule local concentration

The diffusion of QS molecules in the nano-inlets is negligible as advection dominates (see details in the following section). Hence, we stated that QS molecules concentration is null at nano-inlets ($x = 0$ µm). Considering advection together with the confinement compartment axial symmetry at $x = x_{center}$ (Fig. S8), we further assumed that the concentration of QS molecules is maximal at the center of the chamber, leading to the following boundary conditions:

$$\begin{cases} C(0) = 0 \\ \dfrac{\partial C}{\partial x}(x_{center}) = 0 \end{cases}$$

Taking these boundary conditions, we solved Eq. 6 and obtained the following expression, predicting an increase of the local concentration of QS molecules with the number of cells ($N_{total}$) and the distance to the nano-inlets ($x$):

$$C(x, N_{total}) = \frac{s_{eff} N_{total}}{\varepsilon V_{total} \|\langle\vec{v}\rangle_x\|} \left[ \frac{\varepsilon D_{eff}}{\|\langle\vec{v}\rangle_x\|} \, e^{\frac{-x_{center}\|\langle\vec{v}\rangle_x\|}{\varepsilon D_{eff}}} \left(1 - e^{\frac{x\|\langle\vec{v}\rangle_x\|}{\varepsilon D_{eff}}}\right) + x \right] \quad (7)$$

In a regime where advection dominates, one can note that QS molecules concentration varies linearly with the distance from the feeding inlets $x \in [0\,;\, x_{center}]$ (Fig. S6c, insert):

$$C(x, N_{total}) \cong \frac{s_{eff} N_{total}}{\varepsilon V_{total} \|\langle\vec{v}\rangle_x\|} x \quad (7a)$$

#### Inferring the average velocity in the porous medium

##### Hydrodynamic resistance of the nano-inlets

We aim to estimate the average velocity$||\langle\vec{v}\rangle_x||$ in each of the 32 confinement-compartment:

$$||\langle\vec{v}\rangle_x|| = \frac{Q_c}{S_c} \quad (8)$$

The electric-hydraulic analogy (Fig. S9) of the microfluidic device and application of the Kirchhoff's laws give us:

$$\begin{cases} Q_0 = \sum_1^{32} Q_c & (9a) \\ \Delta P_{in} = Q_0 \left( R_{main} + \frac{R_c}{32} \right) & (9b) \end{cases}$$

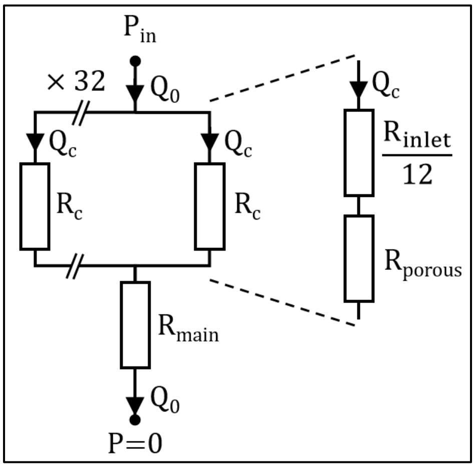


**Figure S9**: Schematic equivalent electronic circuit of the microfluidic device. $P_{in}$ is the applied pressure at the device entrance, $Q_0$ is the flow rate in the main circulating channels (10 µm height) and $R_{main}$ the total hydrodynamic resistance of the main channels. $Q_c$ and $R_c$ are the flow rate and equivalent hydrodynamic resistance of the 32 parallel confinement compartment units. We considered all these units to have a similar resistance over time with a serial of the parallel nano-inlets $R_{inlet}$ and the porous medium (confined proliferating cells) $R_{porous}$.

where, $Q_0$ is the flow rate in the main circulating channels, $Q_c$ is the flow rate within each of 32 parallel confinement compartments and $\Delta P_{in}$ is the pressure applied at the entrance of the microfluidic device. $R_{main}$, $R_c$ are the equivalent hydrodynamic resistance of the main circulating channels and of the 32 confinement compartments respectively. As there are 12 parallel nano-inlets connected to each confinement-compartment, we define $R_c$ as the serial resistance:

$$R_c = \frac{R_{inlet}}{12} + R_{porous} \qquad (10)$$

where, $R_{inlet}$ and $R_{porous}$ are the hydrodynamic resistance of each nano-inlet and the porous medium respectively.

We aim to estimate the average velocity $||\langle \vec{v} \rangle_x||$ in each confinement-compartment:

$$||\langle \vec{v} \rangle_x|| = \frac{Q_c}{S_c} = \frac{Q_0}{32 \times S_c} = \frac{\Delta P_{in}}{32 \times (R_{main} + \frac{R_c}{32}) S_c} \qquad (11)$$

where $S_c$ is the confinement compartment transverse section ($10 \times 40\ \mu m^2$).

The hydrodynamic resistance of a fluidic channel of length L with a rectangular cross section ($h \times w$, $w > h$) can be estimated as:

$$R_{channel} = \frac{12 \eta L}{w h^3 (1 - 0.63 \frac{h}{w})} \qquad (12)$$

As the height of the main circulating channels $h_{main}$ is 10 µm and that of the nano-inlets h is 0.5 µm we get $h_{main}{}^3 \ll h^3$. Since the hydrodynamic resistance of each chamber is at least of the order of magnitude of $\frac{R_{inlet}}{12}$, we neglected the quantity $R_{main}$ and simplified Eq. (11) as:

$$||\langle \vec{v} \rangle_x|| \cong \frac{\Delta P_{in}}{R_c S_c} \qquad (13)$$

**Estimation of the hydrodynamic resistance of the porous medium**

We used the Kozeny-Carman formalism to express the intrinsic permeability of the porous medium ($m^2$):

$$k = \frac{\varepsilon^3}{K_K S_p^2 (1-\varepsilon)^2} \qquad (14)$$

where, $S_p$ ($m^{-1}$) is the specific surface of the particles composing the porous medium ($S_p = 1.13\times10^6$ $m^{-1}$ for uncompressed *Saccharomyces cerevisiae* yeast[1]) and $K_K$ is the Kozeny-Carman empirical factor. Value of this empirical factor is constant for densely packed hard spheres network ($K_K = 5$) but varies in the case of polydispersed and compressible particles like yeasts. Valencia et *al.* estimated $K_K$ values for similar porosity range ($\varepsilon \sim 0.15$ to 0.30) in *Saccharomyces cerevisiae* cells assemblies[4] . To obtain a continuous function over porosity range explored in our case, we fitted $K_K$ values from Valencia et al. with a first order exponential. Starting from $7\times10^{-15}$ $m^2$ at confluency, permeability rapidly decreases with GIP down to $\sim3.5\times10^{-15}$ $m^2$. It then enters in slower decay regime for mechanical pressure values above ~0.2 MPa. We posit that two contributions likely decrease pore radius leading the first sharp decrease: important cellular rearrangements coupled to elevated polydispersity linked to numerous budding events. As cell division rate decreases under elevated pressure (Fig. S6a), densely packed cells tend to accumulated in $G_1$ unbudded state and diminution of pore size is probably mainly modulated by cell deformability.

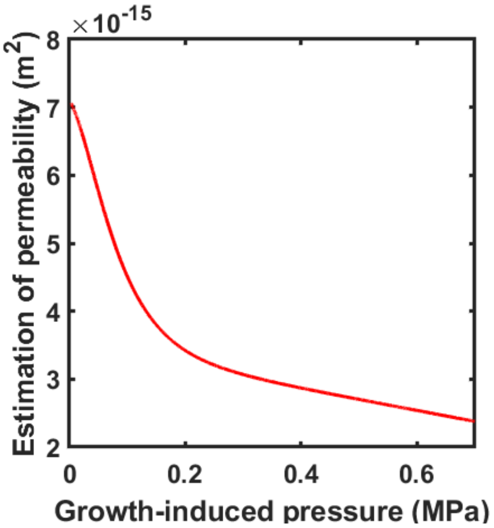

**Figure S10**: Permeability of the porous medium (confinement cells) as function of GIP. Permeability as estimated using the Kozeny-Carman formalism

Having estimated the permeability, we calculated the hydrodynamic resistance of the porous medium:

$$R_{porous} = \frac{\eta L_{porous}}{k S_{porous}} \quad (15)$$

where, η is the fluid dynamic viscosity of the fluid (approximated to water at 30°C, Pa.s); $L_{porous}$ is the total length of the porous medium crossed by the fluid (m) and $S_{porous}$ is its transverse section (m²). We considered that $S_{porous} = S_c$ and $L_{porous} = x_{center}$. As expected, the hydrodynamic resistance of the porous increases with compression, leading to a minor decrease of the average velocity (Fig. S11).

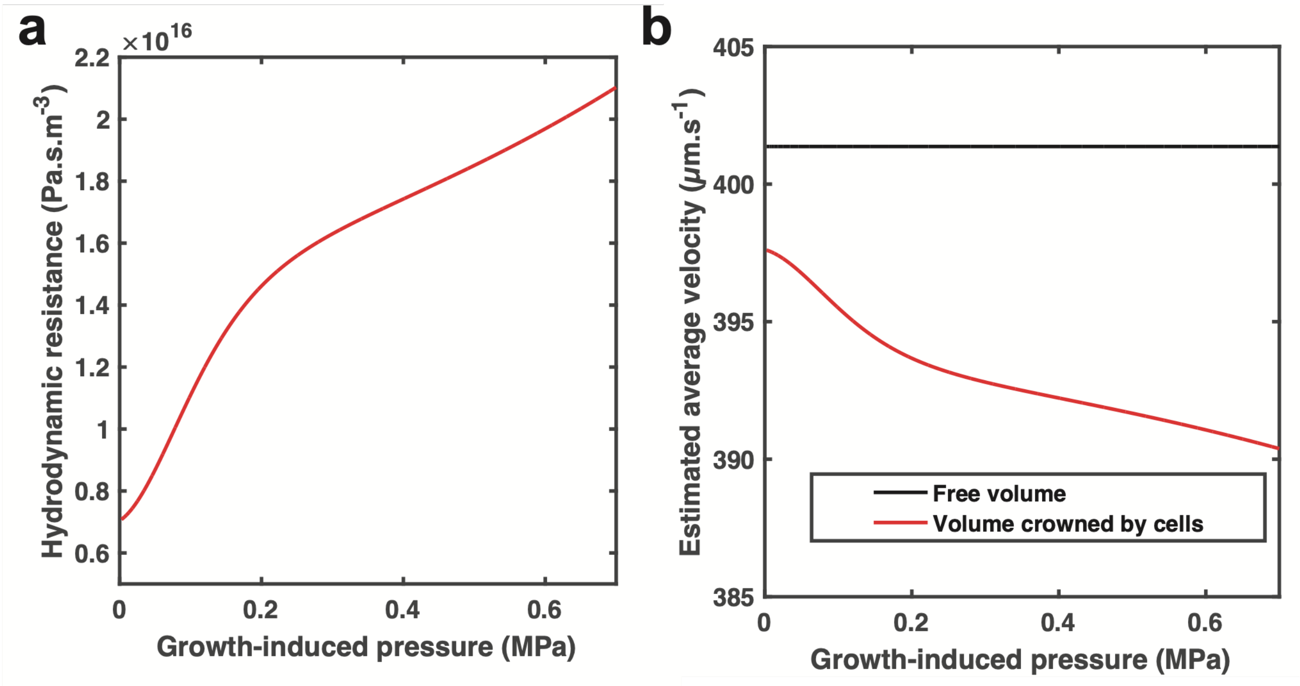


**Figure S11**: **a.** Hydrodynamic resistance of the porous medium (confined cells) increases with growth-induced pressure. **b.** Estimated average flow velocity in one confinement compartment. Flow velocity slightly decreases (3 %) when considering porous medium hydrodynamic resistance.

We obtained $R_{porous} \sim 2 \times 10^{16}$ Pa.s.m$^{-3}$, which is one order of magnitude smaller than the hydrodynamic resistance of the parallel nano-inlets connecting each side of the confinement compartment ($\frac{R_{inlet}}{6} \sim 3.7 \times 10^{17}$ Pa.s.m$^{-3}$). Thus, the average velocity in the porous medium along the half-width of the confinement-compartment can be estimated by (going from 12 inlets on both sides to 6 on one side):

$$\|\langle\vec{v}\rangle_x\| \cong \frac{\Delta P_{in}}{\frac{R_{inlet}}{6} S_{porous}} \qquad (16)$$

**Advection dominates within the nano-inlets**

We estimated the molecular Péclet number ($Pe_{inlet}$) within a nano-inlet to assess whether diffusion or advection dominates. Taking the width of a nano-inlet as its characteristic size $L_i \sim w$, one gets:

$$Pe_{inlet} = \frac{L_i \bar{v}_{inlet}}{D_{mol}} \cong \frac{w^2 h^3 (1 - 0.63\frac{h}{w}) \Delta P_{in}}{12 \eta w h L D_{mol}} \qquad (17)$$

where, w, h and L are the width, height and length of the nano-inlet respectively (L>w>h). We obtained $Pe_{inlet} \sim 72$, suggesting advection dominates in the nano-inlets ($Pe_{inlet} \gg 1$). Hence, one can reasonably assess that the concentration of farnesol is negligible at the nano-inlets (C(x=0) = 0), which justifies the use of Eq. 7a. Thus, the concentration of QS molecules linearly increases between inlet and the center of the chamber.

**Parameter adjustment of the 1-D modeling**

Incorporating the expression of local concentration of QS molecules from Eq. 7a in $\mathbb{P}_{QS}$ we obtain:

$$\mathbb{P}_{QS}(x, N_{total}(t)) = \frac{1}{1 + \frac{S_{eff} N_{total}(t) x}{C_{50} \varepsilon V_{total} \|\langle\vec{v}\rangle_x\|}} = \frac{1}{1 + \frac{x}{x_0}}$$

This is the equation we used to fit the spatial dependence of yeast-to-hyphal transition.

| **Parameter** | **Symbol** | **Units** | **Value** |
|---|---|---|---|
| Confinement compartment length | $l_c$ | m | $4\times10^{-5}$ |
| Confinement compartment width | w | m | $5\times10^{-5}$ |
| Confinement compartment half-width | $x_{center}$ | m | $2.5\times10^{-5}$ |
| Confinement compartment height | $h_c$, $h_{main}$ | m | $1\times10^{-5}$ |
| Confinement compartment transverse cross-section | $S_c$ | $m^2$ | $4\times10^{-10}$ |
| Nano-inlet characteristic length | $L_i$ | m | $1\times10^{-6}$ |
| Nano-inlet length | L | m | $4\times10^{-5}$ |
| Nano-inlet width | w | m | $1\times10^{-6}$ |
| Nano-inlet height | $h_{inlet}$ | m | $5\times10^{-7}$ |
| Hydrodynamic resistance of inlets | $R_{inlet}$ | $Pa.s.m^3$ | $5\times10^{18}$ |
| Length of crossed-flow porous medium | $L_{porous}$ | m | $25\times10^{-6}$ |
| Transverse section of porous medium | $S_{porous}$ | $m^2$ | $4\times10^{-10}$ |
| Specific surface of yeasts | $S_p$ | $m^{-1}$ | $1.13\times10^6$ |
| Total cell monolayer volume | $V_{total}$ | $m^3$ | $1\times10^{-14}$ |
| Dynamic viscosity of culture medium<br>(approximated to water, 30°C) | $\eta$ | Pa.s | $8\times10^{-4}$ |
| QS molecule free diffusion<br>(approximated to glucose 30°C water) | $D_{mol}$ | $m^2.s^{-1}$ | $7.4\times10^{-10}$ |

| Glossary of parameters symbol and units | | |
|---|---|---|
| **Parameter** | **Symbol** | **Units** |
| Applied pressure at device entrance | $\Delta P_{in}$ | mbar |
| Hydrodynamic resistance of main circulating channels | $R_{main}$ | Pa.s.m$^3$ |
| Hydrodynamic resistance of a nano-inlet | $R_{inlet}$ | Pa.s.m$^3$ |
| Peclet number in nano-inlets | $Pe_{inlet}$ | 1 |
| Hydrodynamic resistance of a confinement compartment filled with cells | $R_c$ | Pa.s.m$^3$ |
| Average flow velocity in the porous medium | $\|\langle\vec{v}\rangle_x\|$ | m.s$^{-1}$ |
| Flow rate in confinement compartment | $Q_c$ | m$^3$.s$^{-1}$ |
| Flow rate at microfluidic device output | $Q_0$ | m$^3$.s$^{-1}$ |
| Distance relative to inlets | x | m |
| Total cell number (monolayer) | $N_{total}$ | 1 |
| Total number of cell monolayer at confluency | $\bar{N}_{confluency}$ | 1 |
| Duration for yeast population to reach confluency in confinement compartment | $\bar{t}_{confluency}$ | h |
| Time | t | h |
| Growth-induced pressure | P | MPa |
| Cell division rate | $k_{div}$ | h$^{-1}$ |
| Characteristic pressure of division rate exponential decay | $P_c$ | MPa |
| Cell density | $\rho$ | 1 |
| Total volume of cells | V | m$^3$ |
| Cell packing fraction | $\phi$ | 1 |
| Porosity of the packed cells (porous medium) | $\varepsilon$ | 1 |

| | | |
|---|---|---|
| Hydrodynamic resistance of porous medium | $R_{porous}$ | Pa.s.m$^3$ |
| Kozeny-Carman empirical constant | $K_K$ | 1 |
| Permeability of porous medium | k | m$^2$ |
| Tortuosity of porous medium | T | 1 |
| Typical farnesol diffusion time in the porous medium | $\tau$ | s |
| Effective diffusion of farnesol in porous medium | $D_{eff}$ | m$^2$.s$^{-1}$ |
| Average free medium volume per cell | $V_{elementary}$ | m$^3$ |
| Farnesol local concentration | C | mol.L$^{-1}$ |
| Half inhibiting dose of yeast-to-hypha induction by farnesol | $C_{50}$ | mol.L$^{-1}$ |
| Single-cell effective secretion of farnesol | $s_{eff}$ | mol.s$^{-1}$ |
